# A constitutively active cortical nitrate-uptake cell state underlies nitrogen-use efficiency in perennial ryegrass

**DOI:** 10.64898/2026.10.12.758384

**Authors:** Mallikarjuna Rao Kovi, Precious Guevarra, Sara Thu, Emma Tomine Morvik Kragholm, Akhil Reddy Pashapu, Felix Hernandez Nohr, Igor Ferreira da Costa, Martin Steinkopf, Kristin Håland Gylstrøm, Helga Amdahl, Prabha Manishankar, John Michael Alcantara, Susanne Windju, Manikandan Veerabagu, Muath Alsheikh, Siri Fjellheim, Romulo Sacramento Sobral, Odd Arne Rognli

## Abstract

Perennial ryegrass (*Lolium perenne*) is the most widely sown forage grass of temperate agriculture and a major sink for fertiliser nitrogen, yet the cellular basis of its nitrogen-use efficiency (NUE) is unknown. We built a single-nucleus atlas of 76,684 root nuclei from a high- and a low-NUE genotype under matched nitrogen resupply. High-affinity nitrate uptake is concentrated in one cortical cell state that co-expresses nine NRT2 transporters and all three NAR2 partner proteins, a state that whole-root RNA-seq of the same tissue cannot resolve. Under reduced nitrogen the low-NUE genotype induces this state, whereas the high-NUE genotype does not; it expresses NRT2.3 at a higher baseline and uses different NRT2 paralogs, with one array member nearly silent. Across 44 annual and perennial Pooideae genomes, the same gene families show no life-history difference in copy number, evolutionary rate or selection intensity. Their regulation, however, varies in the same direction as between the two genotypes: the perennial *Brachypodium sylvaticum* expresses the NRT2.3 ortholog at the level that the annual *B. distachyon* reaches only under nitrogen deprivation, and an N-efficient barley genotype does not induce the transporters that an N-sensitive genotype induces. NUE differences are therefore associated with constitutive versus inducible deployment of a conserved toolkit, and the expression and presence/absence variants we identify are candidate breeding markers. The atlas, the first for a forage grass, is released as an open web resource.

## Introduction

Globally, little more than half of the N applied to cropland is recovered in harvested products, and this efficiency has not improved since the 1980s^1^; raising it is central to meeting food demand within planetary limits^2^. Grassland agriculture is a large and inefficient part of this budget. In grazed pasture, ruminants excrete 70-95% of the N they ingest, and of the N deposited in urine patches about 20% is leached, 13% volatilised as ammonia and 2% emitted as nitrous oxide, with only 41% recovered by the sward^3,4^. Management reviews list plant selection for higher nitrogen-use efficiency (NUE) among the few mitigation options that do not reduce output^5,6^, and perennial ryegrass is the most widely sown perennial forage grass of temperate regions^7^.

The genetics of NUE in forage grasses is almost unstudied. Genomic work in perennial ryegrass has concentrated on persistence and winter survival, including population structure under divergent selection for freezing tolerance^8^ and transcriptome contrasts between genotypes, among them resistant and susceptible genotypes of the cultivar Fagerlin infected with pink snow mould^9^ and genotypes of contrasting freezing tolerance^10^; breeding priorities for European grasslands now stress faster genetic gain through genome-enabled selection^11^. Our gene-level understanding of crop NUE comes from annual cereals, where natural alleles of nitrate transporters and their regulators have been identified and deployed: *NRT1.1B* in rice^12^, *OsNRT2.3b*^13^, *TaNPF7.6* in wheat^14^ and the *BRN1* regulator of NLP3 in rice^15^. Perennial grasses use N differently. They maintain root and crown reserves across seasons, remobilise N between tillers and regrowth cycles, and are harvested as vegetative biomass, not as grain. Whether the determinants of NUE found in annuals apply to perennials is not known.

Nitrate acquisition is organised by cell type. In Arabidopsis the NRT1.1 transceptor and the NIN-like proteins (NLPs) trigger a primary nitrate response within minutes^16–18^, high-affinity uptake is carried by NRT2 transporters that require NAR2/NRT3 partner proteins^19^, and single-cell studies show that the outer root cell layers carry the major transcriptional response to soil conditions^20^. Single-cell and single-nucleus atlases of maize, rice and wheat roots have mapped nitrate-responsive genes to cell types^21–24^, a comparison of Brassicaceae root atlases has shown how cell-type programmes diverge between related species^25^, and single-cell multiomics has been applied to the nitrogen response of tomato roots^26^. No such data exist for a perennial grass or a cool-season forage, and no study has compared genotypes that differ in NUE at the level of cell states.

Annual and perennial life histories have evolved repeatedly in grasses, and perennials differ from annuals in allocation and in rates of molecular evolution^27^. Perennial grasses can retain far more applied N than annual cereals^28,29^, which suggests that selection on N economy differs between the two life histories, yet NUE genes have not been compared across annual and perennial lineages. NUE could diverge through changes in the transporters and enzymes themselves, in their copy number or coding sequence, or through changes in where and when a conserved toolkit is expressed; general evidence favours cis-regulatory and duplicate-gene divergence as a common route of phenotypic evolution^30–33^.

Here we built a single-nucleus atlas of whole roots of a high-NUE (Barhoney) and a low-NUE (Fagerlin) genotype of perennial ryegrass, grown as clonal replicates under identical N starvation and resupply, and compared it with bulk RNA-seq of the same tissue, with the N-use gene families of 53 grass genomes spanning annual and perennial Pooideae, and with public root transcriptomes of barley genotypes and of annual and perennial *Brachypodium*. We show that high-affinity nitrate uptake is concentrated in a cortical cell state that whole-root RNA-seq cannot resolve; that the high- and low-NUE genotypes run this state constitutively and inducibly, through different NRT2 paralogs; that the N-use toolkit shows no life-history signature in copy number, evolutionary rate or selection intensity; and that the constitutive-versus-inducible pattern recurs between barley genotypes of contrasting NUE and between an annual and a perennial *Brachypodium*. The atlas is released as an open web resource.

## Results

### Whole-root RNA-seq does not resolve the nitrogen response

We began with bulk RNA-seq of whole roots of the two genotypes and a third cultivar over a resupply time course (full N, 15% and 25% less N; 0 to 48 h; 114 libraries; Fig. 6). Genotype differences were present at every time point and N level (224 to 1,417 differentially expressed genes between Barhoney and Fagerlin per contrast), whereas the N-level contrasts varied widely between time points and genotypes (120 to 1,394 genes) and did not involve the genes expected to carry a nitrogen response: the high-affinity nitrate transporters were barely measurable (NRT2.3 pair ≤ 3 TPM) and the N-uptake and assimilation genes showed no coordinated change with N level. Either the root does not regulate nitrate uptake transcriptionally at this scale, or the response is confined to cells that whole-root RNA averages away. To distinguish the two, we profiled nuclei of the same tissue at the 4 h time point.

### A single-nucleus atlas of the perennial ryegrass root

We profiled nuclei from whole root systems of Barhoney and Fagerlin, two diploid genotypes selected as high- and low-NUE extremes of a two-year field trial of 40 cultivars at two Norwegian sites (Ås and Hamar) under three N regimes. Clonal tillers were N-starved for 7 days and resupplied for 4 h with full N (control) or 25% less N, three plants per group (12 libraries; Fig. 1a). Libraries were sequenced to 91-121 Gb and aligned to the Kyuss_2.0 assembly^34,35^. Cell Ranger called 94,786 nuclei; after ambient-RNA correction (5-15% contamination per library), doublet removal (9-17%) and quality filters, 76,684 nuclei remained, with a median of 889-1,355 genes per nucleus (Supplementary Table 1; Supplementary Fig. 1). About 45% of raw reads derived from rRNA, which halved usable depth but did not affect cell calling, and replicate libraries were highly concordant (within-group pseudobulk Spearman ρ = 0.96-0.99).

**Fig. 1.**
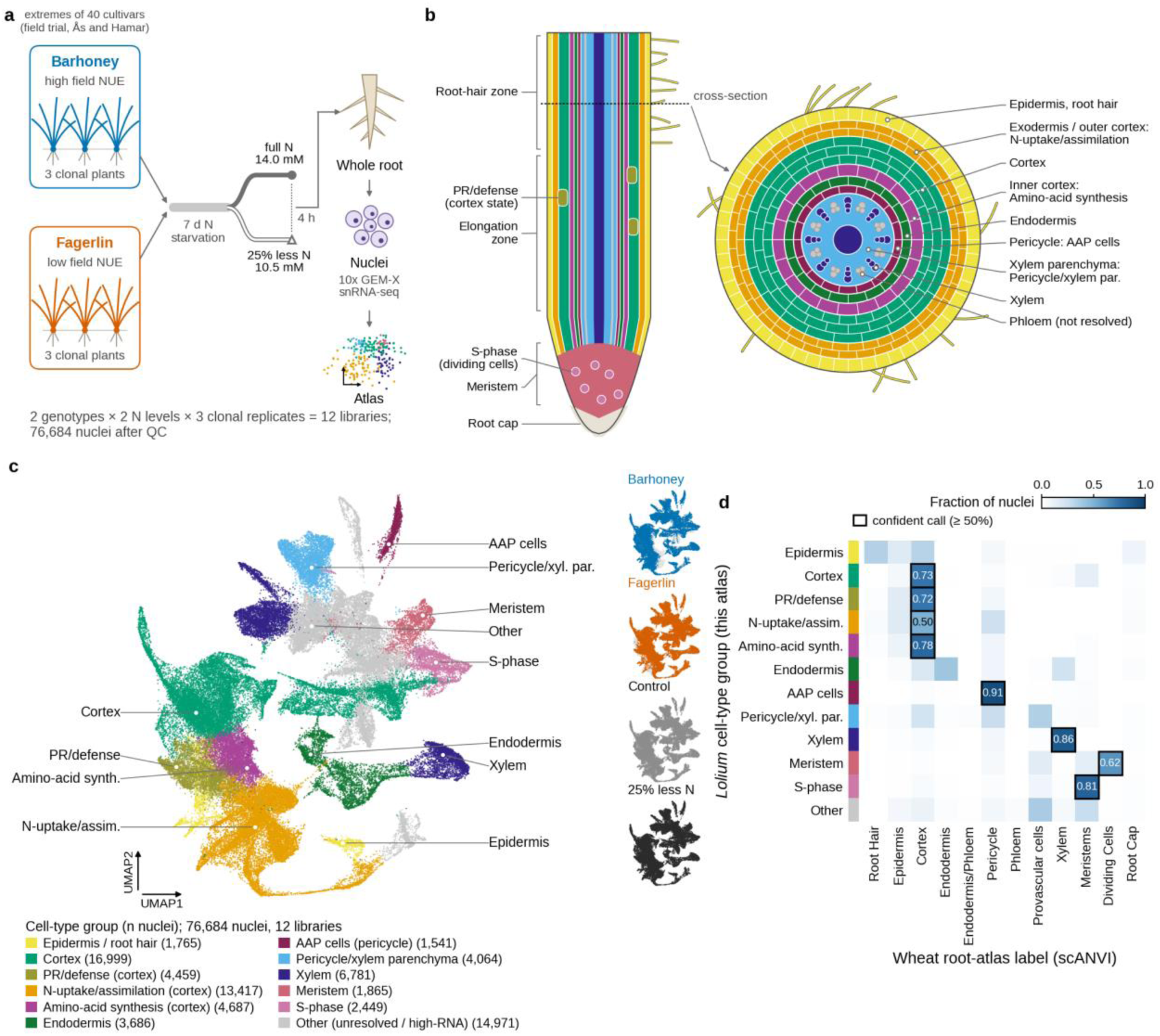
A single-nucleus atlas of the perennial ryegrass root. **a**, Design: two genotypes, the high-NUE (Barhoney) and low-NUE (Fagerlin) extremes of a field trial of 40 cultivars at Ås and Hamar, × two N resupply levels (full N, 14.0 mM nitrate; 25% less N, 10.5 mM) × three clonal replicates; whole roots harvested 4 h after resupply following one week of N starvation. **b**, Root schematic (longitudinal section and mature-zone cross-section) with the tissues coloured by the cell state of the atlas that they correspond to; the radial placement of the two cortical nitrogen states is inferred from label transfer. **c**, UMAP of 76,684 nuclei coloured by cell-type group (number of nuclei per group in the legend; four unresolved or high-RNA clusters merged as Other). N-uptake/assimilation, amino-acid synthesis and PR/defense are functional states within the cortex, and AAP cells a state within the pericycle, named by their dominant programme; their tissue is assigned by label transfer (see panel d). At right, the same embedding coloured by genotype (Barhoney, blue; Fagerlin, orange; both N levels pooled) and by N level (control, grey; 25% less N, black; both genotypes pooled): every cell state contains nuclei of both genotypes and both treatments. **d**, Cross-species label transfer: fraction of nuclei of each cell-type group assigned to each wheat root-atlas cell type by scANVI; rows, the *Lolium* cell-type groups of this atlas, with the colour strip of panel c, each row summing to 1; columns, wheat root-atlas cell types; boxed, confident call (≥ 50%). Per-library QC is in Supplementary Fig. 1.

Integration over library and Leiden clustering gave 35 clusters that mixed across libraries (median mixing entropy 0.94; Supplementary Fig. 2) and resolved cortex (16,999 nuclei), a cortical N-uptake state (13,417), xylem (6,781), an amino-acid-synthesis cortex state (4,687), stele parenchyma (4,064), endodermis (3,686), dividing and meristematic cells (4,314), epidermis and root hairs (1,765) and pericycle (1,541); phloem was not resolved (Fig. 1c; Supplementary Table 2). Marker sets from published root atlases were enriched 12- to 130- fold in the expected clusters. Label transfer from the wheat root atlas^23^ (scANVI and scanpy ingest, 14,414 shared genes) and correlation to wheat cell-type signatures gave concordant identities for meristem, pericycle, trichoblast, xylem and phloem clusters (three of three methods) and for stele and endodermis (two of three), and a maize atlas^21^ agreed at the coarse level. The N-uptake and amino-acid-synthesis states were placed in the cortex, with an epidermal component (Fig. 1b): 50% of uptake-state nuclei were assigned to wheat cortex, 25% to pericycle and 17% to epidermis (Fig. 1d), and 95% to maize cortex.

These states have no counterpart in published grass root atlases. In the wheat and maize root-tip atlases, expression of the N-uptake and assimilation families is only weakly biased to the outer layers (root hair, epidermis and cortex carry 34% of the families’ summed expression in wheat against 25% expected; no enrichment in maize), whereas in our whole-root data 35% falls in the two cortical N states against 13% expected (Supplementary Table 14). Of the Lolium cell-state marker genes with one-to-one orthologs, 40% (of 4,023) peaked in the corresponding cell type of the wheat atlas and 20% (of 1,876) in maize, from 74-78% for meristem and dividing-cell markers in wheat down to 15% for the N-uptake state (1.4-fold over chance, permutation P = 0.01) and 10% in maize (not above chance), while general cortex markers were conserved threefold over chance in both species (Supplementary Table 14c). Cortex identity is therefore shared, whereas the uptake-state programme is not; because the reference atlases are of root tips, in which mature cortex is scarce, tissue maturity and lineage divergence cannot be separated here. To our knowledge this is the first single-nucleus atlas of a cool-season forage grass; the closest precedents are root-tip atlases of annual cereals^21–24^.

### Nitrogen metabolism is partitioned into three cell states

The second-largest population (17.5% of nuclei) co-expressed the high-affinity nitrate uptake machinery: nine of the 17 annotated NRT2 transporters, from the chromosome 3, 4 and 6 arrays and each detected in 26-92% of its nuclei; all three NAR2 partner proteins (88-97%), without which NRT2 transporters are inactive^36^; the ammonium transporter AMT1;1-like (LOC127335544, 88%); the K^+^ channel AKT1 and CIPK21 (Fig. 2a). Nitrate reductase, nitrite reductase, the root GS1 isozymes, GS2 and both GOGATs were expressed here but were as high or higher in the two neighbouring cortical states, so uptake is concentrated in one state whereas reduction and amination are shared across the cortex. We refer to this population as cortical nitrate-uptake cells. It is distinct from the root-hair cluster, which expressed NRT2.3 at lower and AMT2;1 at higher levels. Along a diffusion pseudotime^37^ through the cortical lineage (41,182 nuclei rooted in the meristem), the uptake state is the terminal stage (median rank 0.80, after meristem 0.02, dividing cells 0.08, cortex 0.31 and the asparagine-synthesis state 0.54) and connects to the epidermis rather than back to the cortex (Supplementary Fig. 4; Supplementary Table 19). The uptake module rose along this axis identically in both genotypes and treatments (half-maximal at rank 0.575 in all four groups), and the uptake state made up a similar share of lineage nuclei (30-35%), so the genotypes differ neither in when the programme is switched on during maturation nor in how many cells carry it.

**Fig. 2.**
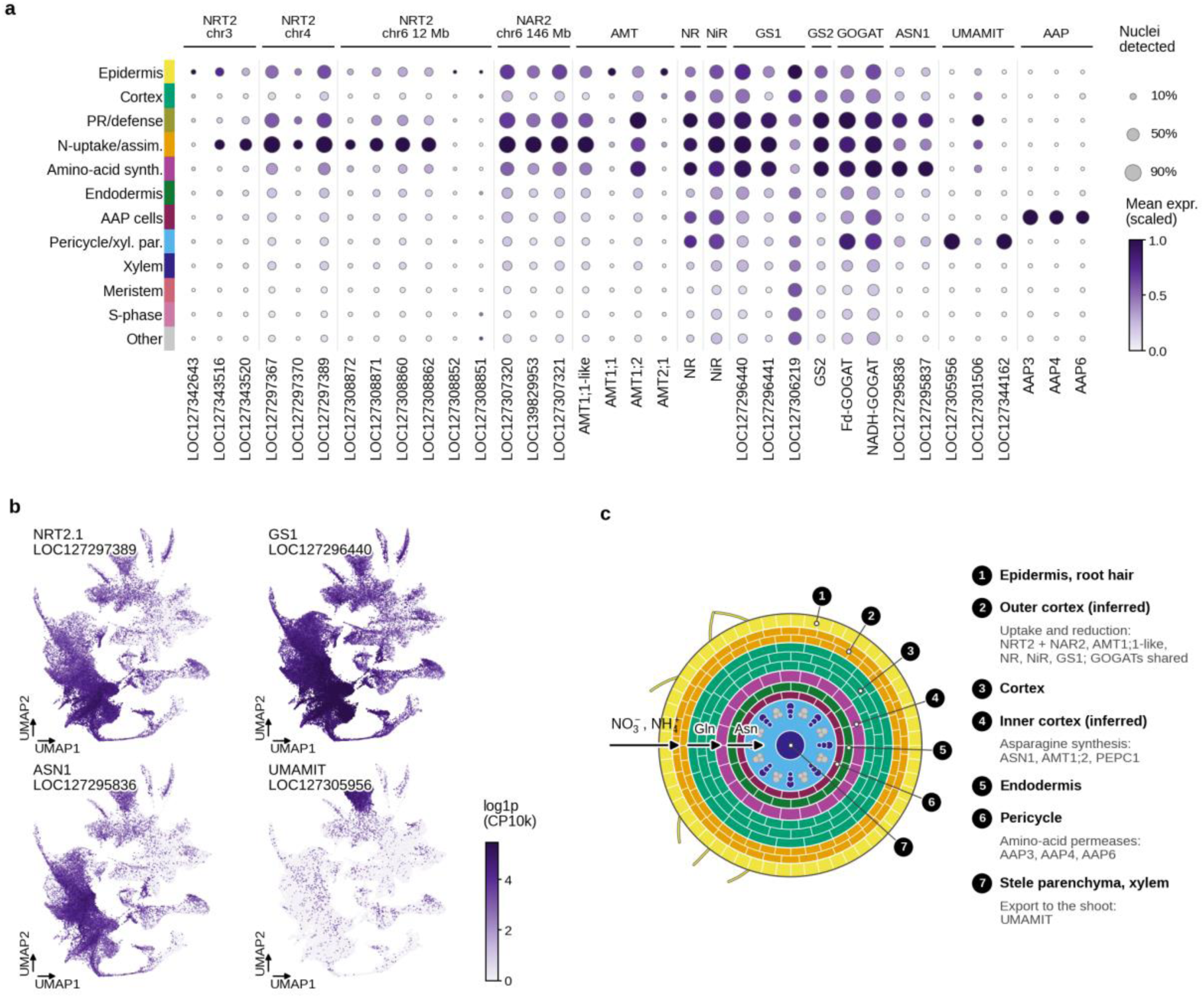
Nitrogen metabolism is partitioned into three cell states. **a**, Dot plot of NRT2 paralogs (by array), NAR2 partners, AMT, NR, NiR, GS1/GS2, GOGAT, ASN, UMAMIT and AAP genes across cell states (dot size, fraction of nuclei detected; colour, mean expression scaled per gene). **b**, Feature plots for NRT2.1, GS1, ASN1 and UMAMIT. **c**, Schematic: outer-cortex uptake and reduction → inner-cortex asparagine synthesis → stele amino-acid loading.

Asparagine synthesis was separate. Both ASN1 copies and the amino-acid transporter AVT6A marked a second cortical state (6% of nuclei), which shared PEPC1, AMT1;2 and NADH-GOGAT with the adjacent PR/defence state. Amino-acid export was separate again: six UMAMIT exporters, PHO1;2, TGAL6 and callose synthase 7 marked stele parenchyma (5%), and AAP3, AAP4 and AAP6 marked pericycle (2%) (Fig. 2a, b). The root therefore partitions nitrate uptake, asparagine formation and amino-acid export into different cell states; their radial order in Fig. 2c is inferred from label transfer. All expressed NRT2 copies sit in the uptake state, with no counterpart of the stele-specific NRT2 that mediates xylem loading in rice^38^. Signalling genes were distributed differently: orthologs of the NRT1.1/NPF6.3 transceptor and the cytokinin-biosynthesis IPT genes scored highest in the pericycle, NLP factors and LBD37/38/39 in pericycle and xylem parenchyma, and NIGT1/HRS1-type GARPs in xylem, whereas the uptake cells scored highest for cytokinin degradation (CKX), ethylene-responsive factors and ABA receptors (Supplementary Fig. 5; Supplementary Table 17). As in Arabidopsis^18^, nitrate acquisition in the cortex is separated from the stele cell types that generate the root-to-shoot cytokinin signal, although these differences in signalling scores are small.

### The nitrogen response is cell-state-specific and almost entirely restricted to the low-NUE genotype

Genotype, not treatment, dominated the transcriptome. Whole-root pseudobulk profiles separated the genotypes on the first principal component (98% of variance), 16,843 genes differed at padj < 0.05 (7,874 higher in Barhoney, 8,969 in Fagerlin), and their direction agreed with bulk RNA-seq for 72-80% of bulk-significant genes (Fig. 3a). By contrast, 25% less N produced no whole-root response at 4 h in either genotype (0 and 7 genes), and cell-type composition did not change with treatment (all q > 0.8), whereas 15 of the 35 clusters differed in abundance between genotypes (q < 0.1; Supplementary Table 11).

**Fig. 3.**
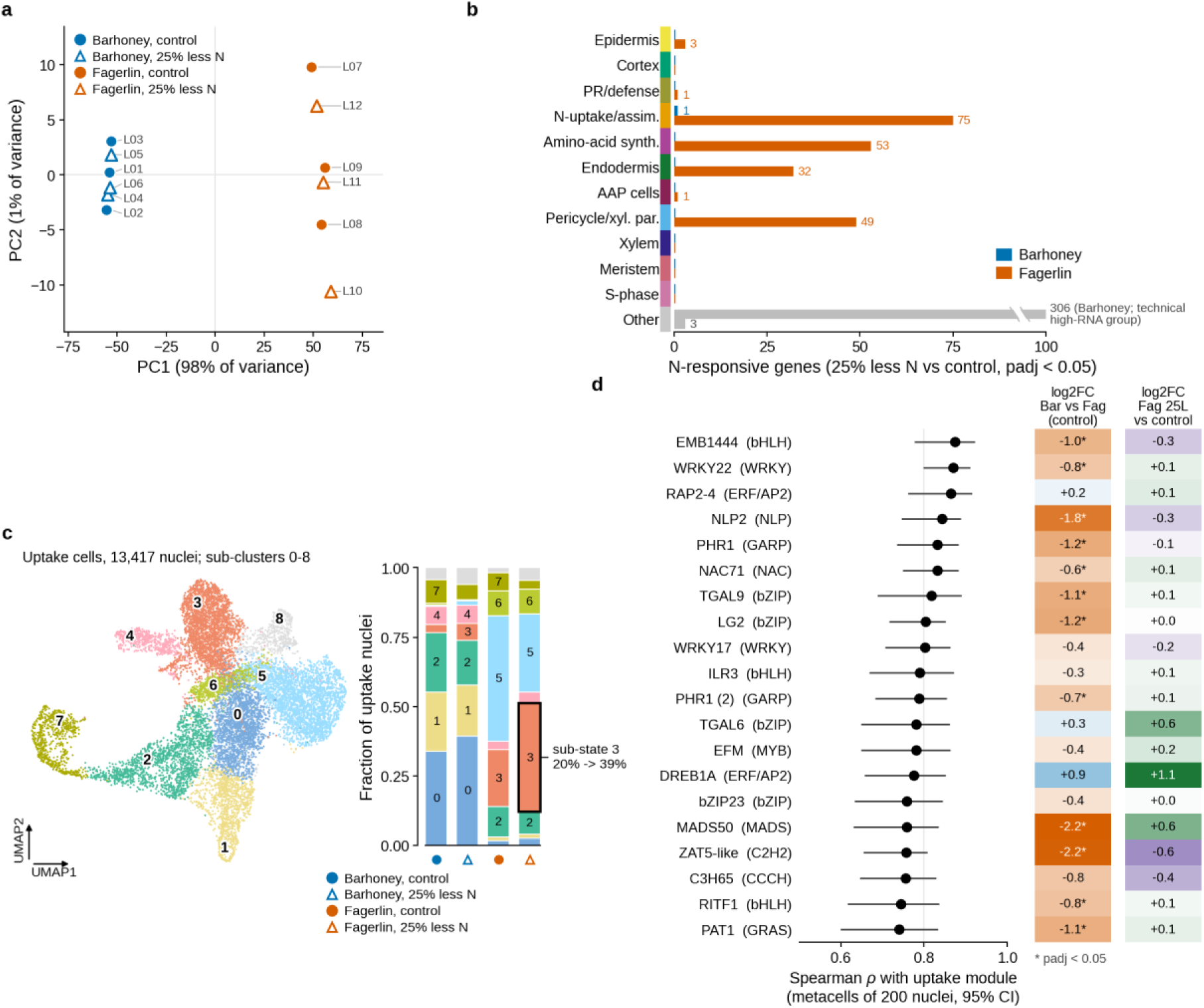
The nitrogen response is cell-state-specific and almost entirely restricted to the low-NUE genotype. **a**, Pseudobulk PCA of the 12 libraries (provider object). **b**, N-responsive genes (25% less N versus full N; DESeq2, adjusted P < 0.05) per cell type and genotype; in the resolved cell states Barhoney has one responsive gene, in the uptake state (the 306 Barhoney genes in the merged unresolved group come from a technical high-RNA cluster). **c**, Uptake-cell sub-clusters by genotype and treatment (share of each genotype’s uptake nuclei), with the induced sub-state and its markers. **d**, Transcription-factor correlation with the uptake module and genotype behaviour.

The N response was present in specific cells and almost only in the low-NUE genotype. Per-cell-type pseudobulk analysis found 75 genes responding to 25% less N in Fagerlin uptake cells (51 up, 24 down), 53 in amino-acid-synthesis cells, 49 in stele parenchyma and 32 in endodermis, against one gene in Barhoney uptake cells (a glucan endo-1,3-β-glucosidase) and none in any other resolved Barhoney cell state (Fig. 3b). Fagerlin contributed 8,102 uptake nuclei against 5,315 for Barhoney, so part of this asymmetry may be power. In Fagerlin uptake cells the response centred on the uptake machinery: both NRT2.3 copies (LOC127343516, LOC127343520; log_2_ fold change +1.15 and +1.13, padj 0.017) and AMT2;1 (+1.33) were induced, alanine aminotransferase 2 (−1.44) and ATG8C (−0.88) were repressed, and “nitrate transmembrane transporter activity” was the top enriched term; hypoxia- and stress-responsive genes (two ADH3 copies, a plant cysteine oxidase) responded as well. The NRT2.3 induction was robust to leaving out any single library and to ambient correction, which removed at most 3% of counts for these genes.

Sub-clustering resolved an induced state. One sub-cluster rose from 20% to 39% of Fagerlin uptake nuclei under 25% less N (676 versus 1,877 nuclei; every 25%-less-N library above every control library) and was marked by the chromosome-6 NRT2.1 copy LOC127308872, a heavy-metal-associated protein, the aquaporin TIP4-3, the clock gene CCA1 and a wall-associated kinase (Fig. 3c). The same sub-state was present in Barhoney at low abundance and shifted in the same proportion (2.8% to 5.8% of its uptake nuclei), so induction is not absent in the high-NUE genotype but an order of magnitude smaller. RNA velocity (10.8% unspliced molecules) showed no treatment-dependent shift in latent time within any state (Supplementary Fig. 3), consistent with induction within a stable cell state, without a transition between states.

Among 333 transcription factors detected in at least 2% of uptake nuclei, those most correlated with the uptake module across metacells included NLP2 (ρ = 0.84), two PHR1-like GARP factors, the TGA-class bZIPs TGAL9 and TGAL6, WRKY22, RAP2-4 and a bHLH, and LBD37 was the third-best correlate of the induced gene set (Fig. 3d; Supplementary Table 4); NLP, LBD and TGA factors belong to the nitrate-signalling network defined in Arabidopsis^17,18,39,40^. These are expression correlates, not demonstrated regulators, and most were higher in Fagerlin at control (NLP2 3.5-fold; TGAL9 and the PHR1-like factor 2.2-fold), the genotype with lower target expression, so the higher baseline of the uptake genes in Barhoney is not explained by more of these factors. A motif-based test of direct regulation was uninformative, because the TGACG and W-box motifs occur in 59-74% of 1kb Lolium promoters.

### The two genotypes deploy different NRT2 paralogs in the same cells

Within the uptake cells the genotypes expressed the NRT2 family to a similar extent but used different members of it (Fig. 4a, b; Supplementary Table 3). Barhoney expressed the chromosome-4 member LOC127297370 2.4-fold and NRT2.3 (LOC127343516) 2.0-fold higher (padj 4 × 10^−5^ and 0.005), together with cytosolic GS1 (2.4-fold, padj 9 × 10^−5^). Fagerlin expressed the chromosome-6 member LOC127308862 6.3-fold (padj 1 × 10^−21^) and both GOGATs 2.7-2.8-fold higher, and a second chromosome-6 member, LOC127308872, was expressed in every Fagerlin library (72-239 CPM in whole-root pseudobulk) and essentially silent in every Barhoney library (0-0.5 CPM). Unique-read coverage showed that the locus is present in Barhoney but transcribed at 1% of the Fagerlin level, so the allele is inactive or divergent, not deleted (Supplementary Table 18). The genes Fagerlin induced under 25% less N were not induced in Barhoney; NRT2.3 was already 2-fold higher in Barhoney at control, whereas AMT2;1 was expressed equally at control (Fig. 4b). The genotypes thus differ in baseline and inducibility for NRT2.3 and in inducibility alone for AMT2;1. Both arrays were detected in 87-98% of uptake nuclei of both genotypes (Supplementary Table 16), and the NRT2.3 pair was detected in 36-82% of Barhoney uptake nuclei across sub-states but, in Fagerlin, in 78% of the induced sub-state and 26-28% of two large basal sub-states. The 17 NRT2 loci of Kyuss_2.0 lie mainly in clusters (seven within 0.4 Mb on chromosome 6, three on chromosome 4, the NRT2.3 pair on chromosome 3; Supplementary Table 12), and closest paralogs are 87-96% identical, so these paralog-level differences rest on uniquely mapped reads and need paralog-specific assays for confirmation.

**Fig. 4.**
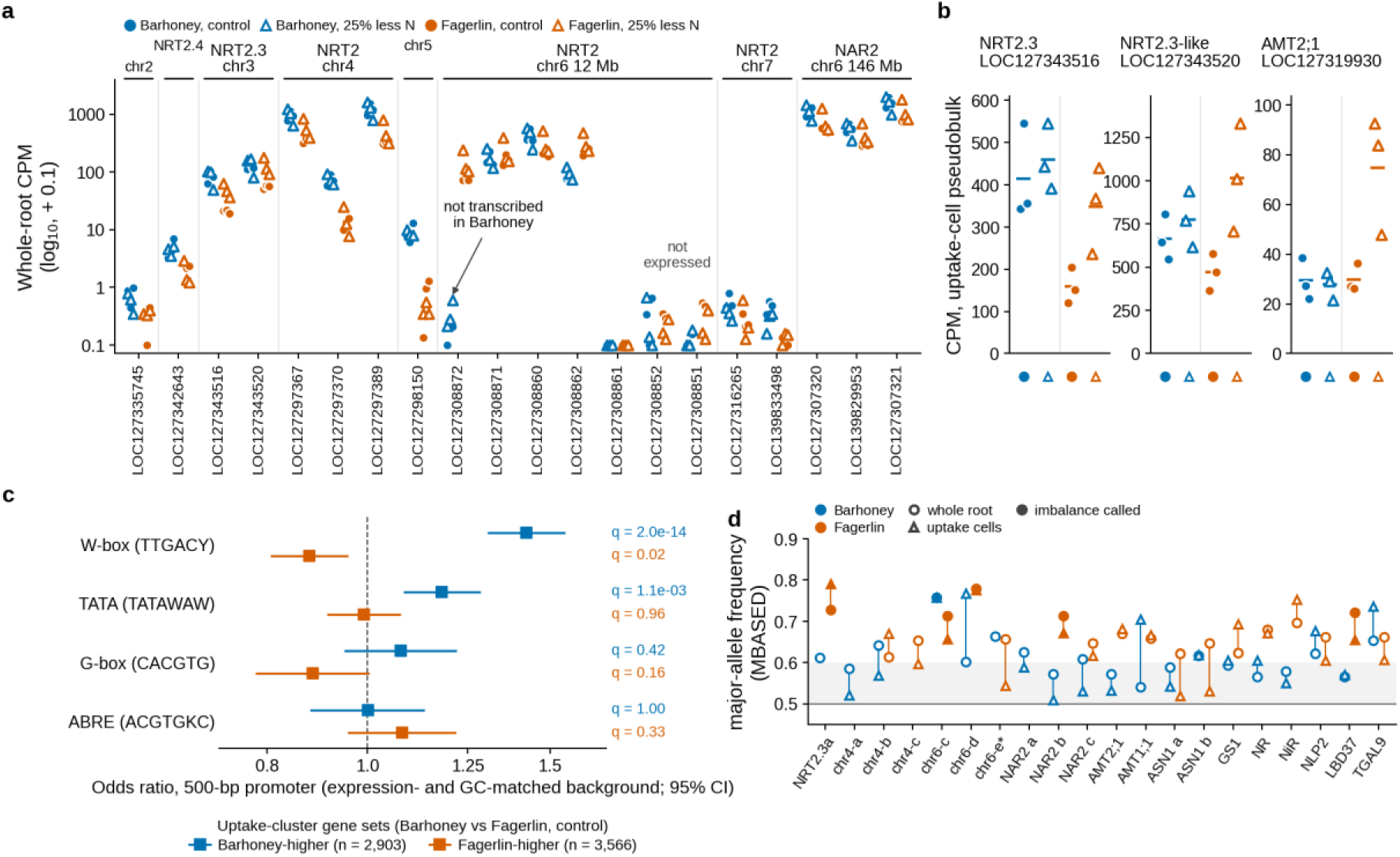
The genotypes deploy different NRT2 paralogs. **a**, All 17 annotated NRT2 genes and the three NAR2 genes: whole-root pseudobulk CPM per library, grouped by chromosomal array; the chromosome-6 member LOC127308872 is expressed in Fagerlin and essentially silent in Barhoney (≤ 0.5 CPM). **b**, NRT2.3, NRT2.3-like and AMT2;1 in uptake-state nuclei (per-library pseudobulk CPM; bars, group means): NRT2.3 higher in Barhoney at control and induced in Fagerlin only; AMT2;1 at the same control level in both and induced in Fagerlin only. **c**, Proximal W-box, TATA, G-box and ABRE elements (500 bp upstream) in genes higher in Barhoney (n = 2,903) or higher in Fagerlin (n = 3,566) in uptake-cluster nuclei at control (provider cluster 0): odds ratio against expression- and GC-matched background genes with 95% confidence interval and BH-adjusted q. **d**, Allele-specific expression at 24 focal genes: MBASED major-allele frequency over each clone’s heterozygous SNPs (uniquely mapped reads; SNPs whose alternative allele occurs elsewhere in the genome removed; counts pooled over six libraries; unphased mode with simulation calibration) in whole roots (circles), Barhoney (blue) and Fagerlin (vermilion); filled, MBASED call (BH-adjusted P < 0.05, frequency ≥ 0.7); triangles, the leave-one-library-out major-allele fraction in uptake-state nuclei (≥100 reads); grey band, 0.5-0.6. chr4/chr6, NRT2.1 array members; chr6-e*, the member inactive in Barhoney (whole-root value only, 104 reads). Supplementary Table 13.

**Fig. 5.**
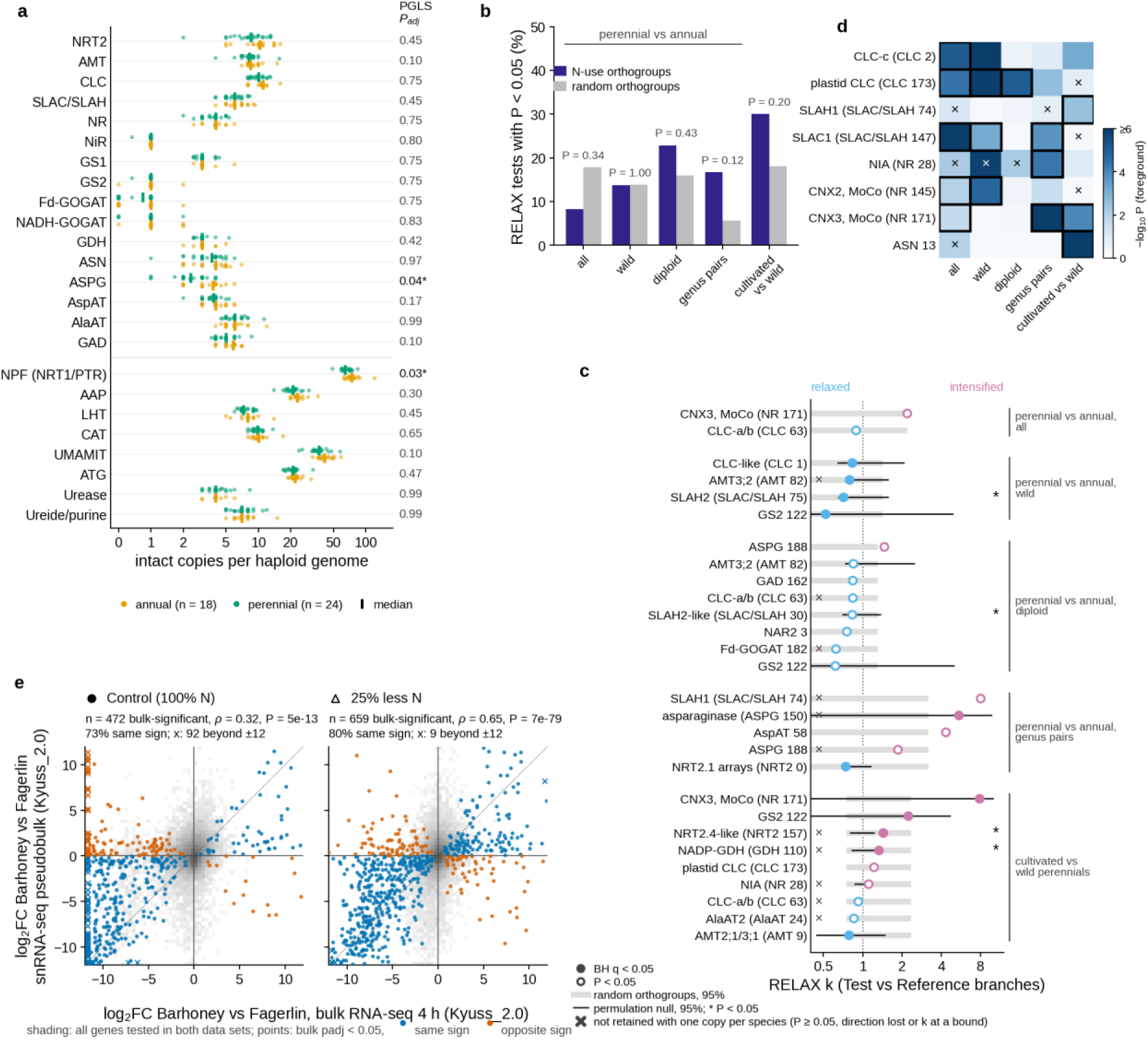
Comparative genomics and bulk consistency. **a**, Intact gene copies per haploid genome (count divided by the ploidy multiplier; zero counts drawn at 0.5) for the 24 families in 42 Pooideae genomes with curated ploidy, annual (orange, n = 18) and perennial (green, n = 24); vertical bars, group medians; right column, BH-adjusted P of the life-history term in a phylogenetic generalised least-squares model with ploidy level and assembly level as covariates (asterisk, P < 0.05). **b**, Fraction of orthogroup × contrast tests with a RELAX shift in selection intensity at P < 0.05, for the N-use orthogroups (indigo; 24-35 interpretable tests per contrast) and for 100 random single-copy orthogroups run through the identical labelling and tests (grey; 70-79 tests); Fisher’s exact P above each pair; after adjustment for the number of sequences per test (logistic regression) the N-use odds ratio is 1.06 (P = 0.85) over all contrasts and 5.9 (P = 0.07) within genera (Supplementary Table 7e). Contrasts: perennial versus annual over all trait-coded genomes, wild genomes only, diploids only and within the five genera containing both life histories; cultivated versus wild perennials. **c**, The 28 interpretable N-use tests with RELAX P < 0.05 (replicate-combined; k of the best-likelihood replicate): selection-intensity parameter k of the Test branches (perennial, or cultivated in the domestication contrast) relative to the Reference branches, k < 1 relaxed (sky blue) and k > 1 intensified (purple); filled, BH q < 0.05 within the contrast; grey bars, central 95% of k over the interpretable random orthogroups of the same contrast; black bars, central 95% of k over the species-tree permulations of the test (42-49 per test; 89 for SLAH2), with an asterisk where permulation P < 0.05; labels give the gene identity of the *L. perenne* members and the family and orthogroup number used in Supplementary Table 7; ×, not retained when each species is represented by one copy (P ≥ 0.05, direction lost or k at a bound). **d**, BUSTED-PH for the orthogroups with at least one contrast passing the three-test rule: shading, −log_10_ P for episodic diversifying selection on the foreground branches; bold outline, foreground significant with no background signal and a foreground distribution different from the background; ×, background branches also significant. CNX2 and CNX3 are molybdenum-cofactor genes mined with the nitrate-reductase seeds. **e**, Barhoney-versus-Fagerlin log_2_ fold changes in bulk RNA-seq at 4 h (x) and in snRNA-seq pseudobulk (y), both quantified on Kyuss_2.0, under full N (left) and 25% less N (right). Grey shading, density of all genes tested in both data sets; points, genes significant in bulk (adjusted P < 0.05), blue where the sign agrees between data sets and vermilion where it does not; crosses, genes beyond the axis limits. Spearman ρ, P and the fraction with the same sign are given above each panel.

**Fig. 6.**
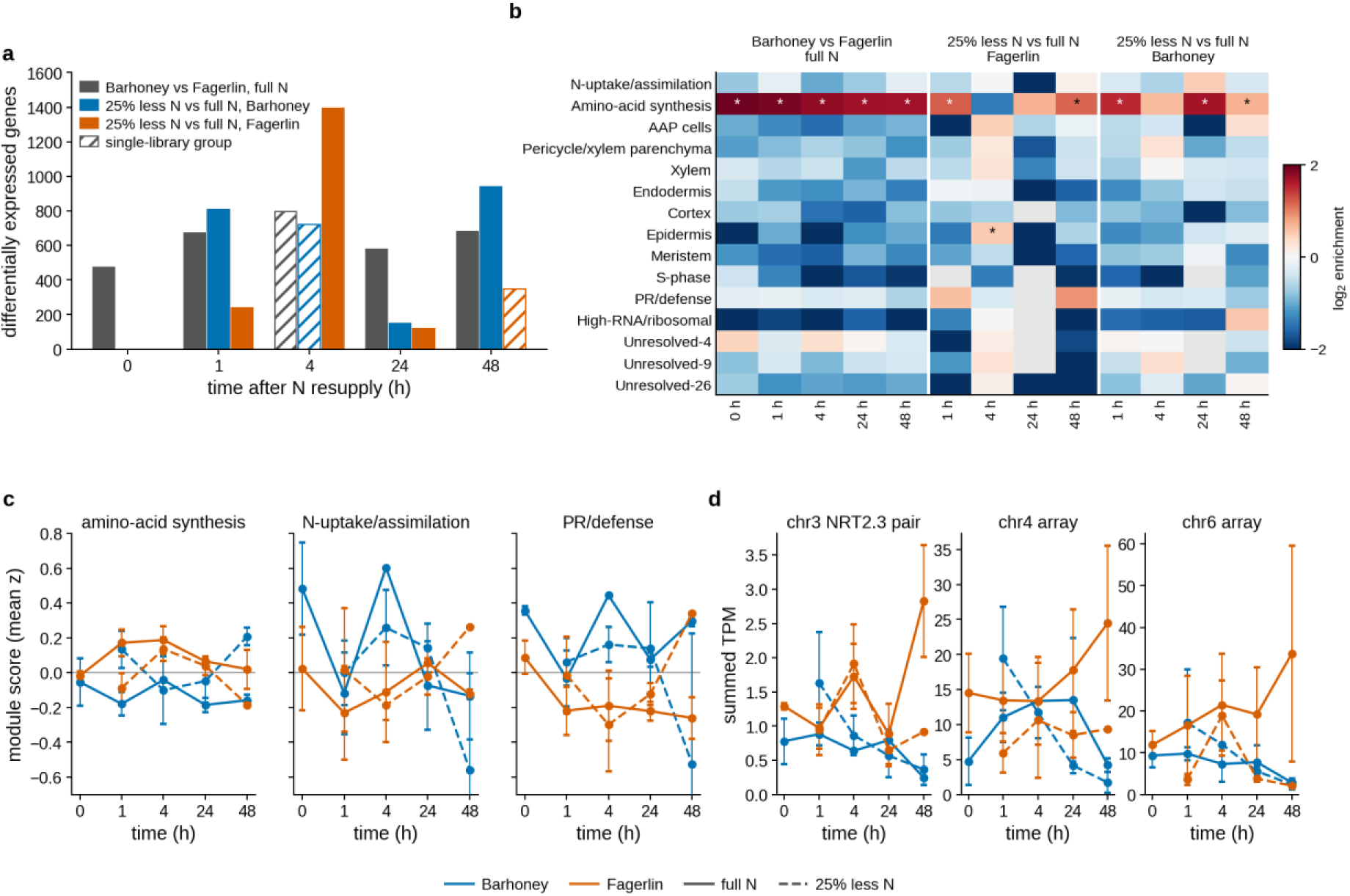
The bulk RNA-seq time course read through the atlas. **a**, Differentially expressed genes (DESeq2, adjusted P < 0.05) at 0, 1, 4, 24 and 48 h after N resupply for Barhoney versus Fagerlin at full N (grey) and for 25% less N versus full N within Barhoney (blue) and Fagerlin (vermilion); hatched bars, contrasts in which one group contains a single library. **b**, Cell-state attribution of each set of differentially expressed genes: log_2_ enrichment of the genes specific to each snRNA-seq cell state (tau ≥ 0.6, ≥ 5 CPM; 401-1,595 genes per state) among the set relative to all expressed genes; asterisks, BH-adjusted one-sided Fisher P < 0.05; grey, fewer than three state-specific genes expected. **c**, Cell-state module scores in bulk (mean z-score of log_2_ TPM over the module genes; mean ± s.e.m. of libraries) over time for the amino-acid-synthesis, N-uptake/assimilation and PR/defence states, by genotype (colour) and N level (solid, full N; dashed, 25% less N). **d**, Summed TPM of the chromosome-3 NRT2.3 pair, the chromosome-4 NRT2 array and the chromosome-6 NRT2 array in bulk over time (mean ± s.e.m.), same encoding as panel c.

Promoter architecture separated the two gene sets. Among the 2,903 genes higher in Barhoney uptake-cluster nuclei at control, proximal W-box elements (TTGACY within 500 bp of the transcription start) were more frequent than in expression and GC-matched background genes (42% versus 34% of promoters; odds ratio 1.42, q = 2 × 10^−14^), as were TATA boxes (odds ratio 1.18, q = 0.001), and the W-box was depleted among the 3,566 Fagerlin-higher genes (odds ratio 0.88, q = 0.02); G-box and ABRE elements were not enriched (Fig. 4c; Supplementary Table 8). Genes higher in the high-NUE genotype are thus modestly enriched for elements of inducible promoters, in the shared Kyuss reference sequence.

We then tested for allelic imbalance within each clone with MBASED^41^ (unphased mode; call rule BH-adjusted P < 0.05 and major-allele frequency ≥ 0.7), with a leave-one-library-out check of library-level reproducibility (Methods). Genome-wide, 6% of Barhoney and 20% of Fagerlin testable genes met the call (803 of 13,089 and 3,330 of 16,341); the higher rate in Fagerlin also reflects its greater divergence from the Kyuss reference, so genome-wide estimates are descriptive (Supplementary Table 13). After removing sites whose alternative allele occurs elsewhere in the reference, MBASED called imbalance at 2 of 21 testable focal N genes in Barhoney and 6 of 19 in Fagerlin (Fig. 4d). NRT2.3 (LOC127343516) was imbalanced in Fagerlin (major-allele frequency 0.73, replicated in five of six libraries) and not in Barhoney (0.61), whereas AMT2;1 was balanced in both (0.57 and 0.67); imbalance at two chromosome-6 array members in both genotypes (0.71-0.78) is as consistent with residual cross-mapping as with cis variation. Imbalance at NRT2.3 only in the low-expressing genotype is what a down-regulating cis variant on one Fagerlin allele would produce, but between-genotype cis and trans effects can only be separated in a Barhoney × Fagerlin F1.

Genome-wide, 508 genes were expressed in one genotype and not the other (395 Fagerlin-only, 113 Barhoney-only; 320 of them lncRNAs). Unique-read coverage classified 384 as expression on/off switches (transcribed in both genotypes but at ≤ 5% in one) and 22 protein-coding genes as lacking uniquely mapped reads in one genotype, six with essentially no reads in gene body or flanks (deletion-like) and 16 with covered flanks (Supplementary Table 5).

The same configuration appears in public data from one barley genotype pair. In root RNA-seq of genotypes described by its authors as N-efficient (W26) and N-sensitive (W20) (N-use efficiency after 21 days of low N 87.5% and 61.7%^42^; sufficient versus low N at 3 and 18 days and resupply at 21 days, three replicates), the N-sensitive genotype induced the orthologs of our uptake-cell genes under low N: the chromosome 3H NRT2 (HvNRT2.5, the reciprocal best hit of Lolium NRT2.3) 8.7-fold at 18 d (P = 8 × 10^−5^), three chromosome-6H NRT2 members 9- to 21-fold at 3 d and the AMT1;1 ortholog 4.3-fold at 18 d (P = 1 × 10^−4^). The N-efficient genotype induced none of them (log_2_ −4.4 to +0.5; P > 0.1) and expressed the 3H NRT2 at a higher level under sufficient N, although not significantly at n = 3 (geometric means 78 versus 4.8 TPM; P = 0.25) (Fig. 7b; Supplementary Fig. 7; Supplementary Table 20). Barley whole roots carry NAR2 and NRT2 transcripts at 10^2^-10^3^ TPM, enough to read without nuclei. This is one genotype pair, but it shows that the pattern is not confined to ryegrass. In three wheat experiments without NUE contrasts, NRT2 family responses to low N had either sign, and in no whole-root data set, ours included, was the orthologous uptake-state signature (39-44 genes) induced as a module (Supplementary Table 20).

**Fig. 7.**
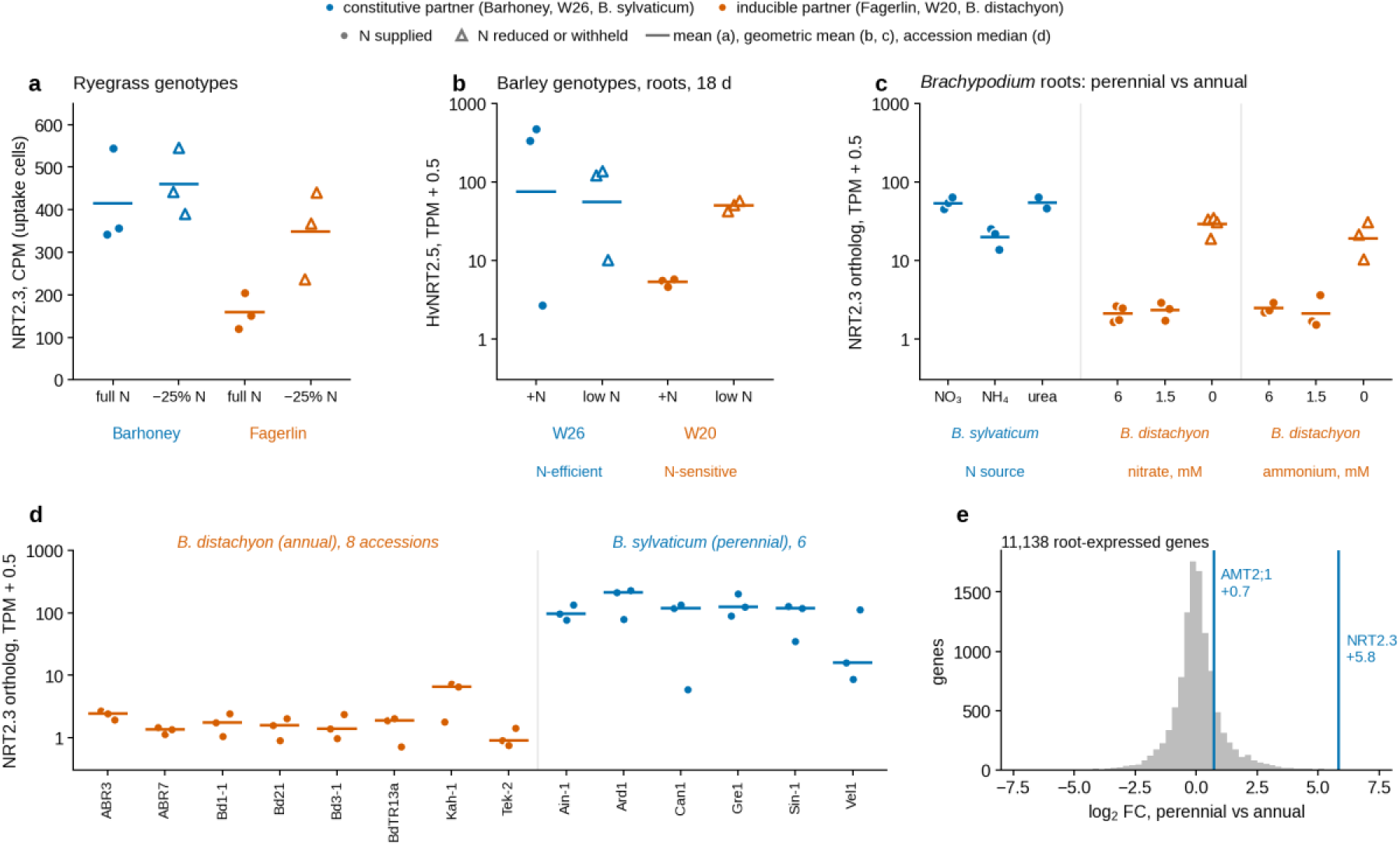
The constitutive-versus-inducible configuration of NRT2.3 recurs between genotypes and between an annual and a perennial grass. Blue, the constitutive partner of each pair (high-NUE Barhoney, N-efficient barley W26, perennial *B. sylvaticum*); vermilion, the inducible partner (low-NUE Fagerlin, N-sensitive barley W20, annual *B. distachyon*); filled circles, N supplied; open triangles, N reduced or withheld; points, libraries or roots; bars, means in panel a, geometric means in panels b and c, and accession medians in panel d. **a**, NRT2.3 (LOC127343516) in uptake-state nuclei of perennial ryegrass, per-library pseudobulk CPM, full N versus 25% less N (as in Fig. 4b). **b**, HvNRT2.5, the reciprocal best hit of *Lolium* NRT2.3, in barley roots at 18 d under sufficient (+N) or low N (public data PRJNA896249; n = 3). **c**, The *Brachypodium* NRT2.3 ortholog in roots of perennial *B. sylvaticum* Ain-1 grown with nitrate, ammonium or urea as N source (PRJNA411649) and of annual *B. distachyon* Bd21-3 at 6, 1.5 and 0 mM nitrate or ammonium (PRJNA1348082). **d**, The same gene per root in one greenhouse batch (JGI R18) of 8 annual *B. distachyon* and 6 perennial *B. sylvaticum* accessions grown together in fertilised soil. **e**, Genome-wide distribution of the perennial-versus-annual log_2_ fold change in that batch (11,138 genes; limma-voom, accession as random block), with NRT2.3 and AMT2;1 marked. TPM, transcripts per million (+ 0.5 on log scales). Full gene sets and the second *Brachypodium* batch are in Supplementary Figs. 7 and 10.

### Nitrogen-use genes show no consistent life-history signal in sequence

We next asked whether the N-use toolkit itself has evolved with life history. We analysed 25 N-use and associated families (uptake, assimilation, remobilisation, autophagy and urea metabolism) across 53 genomes (48 Pooideae, four other grasses and *Arabidopsis*), with the Pooideae core arranged as a 2 × 2 of life history (annual or perennial) and cultivation (wild or cultivated) and both life histories represented within *Alopecurus*, *Brachypodium*, *Hordeum*, *Lolium* and *Poa* (Supplementary Table 6; Supplementary Fig. 6). Gene models were mined uniformly with miniprot^43^, intact loci were counted, and orthogroups were built with OrthoFinder^44^ on a species tree from 87,033 amino-acid sites (IQ-TREE 2^45–47^). Perenniality is ancestral in Pooideae^48,49^, and the panel contains 10 to 17 independent origins of annuality depending on how reversals are weighted. Annual lineages are expected to evolve faster genome-wide for reasons of generation time and population size^50^, so each test asks whether an N-use orthogroup deviates from 100 random single-copy orthogroups run through the identical pipeline, or from species-tree permulations; the null is never one of no change (Methods; Supplementary Note 1).

Copy number did not track life history robustly. Phylogenetic generalised least squares with Pagel’s λ^51^, ploidy and assembly level as covariates (42 genomes; 18 annual, 24 perennial) found perennials to carry slightly fewer NPF (NRT1/PTR) and asparaginase loci (β = −0.14 and −0.31 on log counts; BH-adjusted P = 0.03 and 0.04; Fig. 5a; Supplementary Table 7a), but neither was significant within diploids (BH-adjusted P = 0.20 and 0.63) and no family differed consistently within the five mixed genera (sign tests, P ≥ 0.38; Supplementary Table 7b). NRT2 forms a clustered repertoire in *Lolium* (11 intact loci in *L. perenne*, eight of them in the chromosome-4 and -6 arrays, and 12 and 14 in the other two diploid *Lolium* genomes, against a median of 10, interquartile range 8-11, across 29 diploid Pooideae). Relative evolutionary rates (RERconverge^52^, 500 permulations) did not track life history either: 25 of 236 core tests were nominally significant (2-16% per contrast, against 7-11% of the 148 non-core orthogroups; Fisher’s exact P ≥ 0.27) and 9 at permulation P < 0.05 (3.8%, about the chance rate). The only test at BH q < 0.1, a slower NADP-glutamate dehydrogenase in perennials (q = 0.016, permulation P = 0.008), concerns an orthogroup whose selection result depends on paralog structure (below) (Supplementary Table 7f). Presence of 15 consensus promoter elements in 15,272 upstream regions shifted with life history no more often in N-use orthogroups than in the background (1-4% of tests at permulation P < 0.05 in both sets; none at q < 0.1; Supplementary Table 7h).

Shifts in the intensity of purifying selection were no more common in N-use genes than in random genes. We tested the 50 orthogroups of the 17 core uptake and assimilation families with RELAX^53^ and BUSTED-PH^54,55^ in five contrasts (perennial versus annual over all 44 trait-coded genomes, wild genomes only, diploids only and within the five mixed genera; cultivated versus wild perennials; 248 tests; Supplementary Table 7c). In every contrast the fraction of shifts at P < 0.05 (8-30%) was comparable to that of the random orthogroups (6-18%; Fisher’s exact P ≥ 0.12; Fig. 5b), and with alignment size in a logistic model the N-use orthogroups were no more likely to shift than random genes (adjusted odds ratio 1.06, P = 0.85; within genera 5.9, P = 0.07; Supplementary Table 7e). Of 148 tests with an interpretable selection-intensity parameter k, 28 were nominally significant (Fig. 5c). Replicate fits reproduced 25; aBSREL and Contrast-FEL found no evidence of adaptive episodes within them; 11 lost significance, direction or interpretability when each species was represented by one copy, nine of them with duplication nodes in the Test partition (Supplementary Table 7g); and of the robust shifts tested against 42-49 species-tree permulations, GS2 in three contrasts (P = 0.40-0.86), CNX3 (0.56) and AMT2;1/3;1 (0.26) under domestication, the NRT2.1 arrays within genera (0.08), AMT3;2 among diploids (0.34) and a CLC-like channel in wild perennials (0.32) were no more extreme than chance (Supplementary Table 7i). Four tests that had already failed an earlier step also had permulation P < 0.05 (0.02-0.046) and are not retained. The direction of the nominal shifts was set mostly by the contrast, not by the gene, and random orthogroups that shift are relaxed in perennials in a similar proportion (78%) to the N-use set (68%). Trait-associated episodic selection (BUSTED-PH; 15 tests) was found mainly in anion channels (SLAC1, a plastid CLC, CLC-c, SLAH1) and in the molybdenum-cofactor genes CNX2 and CNX3 (Fig. 5d; Supplementary Table 7d), not in the transporters and enzymes that differ between the two ryegrass genotypes.

One test passed every step. The orthogroup SLAC/SLAH 75 contains a single *L. perenne* gene, LOC127296474, annotated as SLAH2, the nitrate-selective S-type anion channel^56^. Purifying selection on it is relaxed in wild perennials relative to wild annuals (k = 0.71; replicates 0.63, 0.63 and 0.71; q = 0.015), the shift persists with one copy per species (k = 0.86, P = 0.03), and it was more extreme than all but 2 of 89 species-tree permulations (P = 0.033). It appears in no other perennial-versus-annual contrast and shows no adaptive episode. In the atlas the gene is expressed mainly in endodermis and cortex, and endodermal expression is 6.9-fold higher in the high-NUE genotype under full N (log_2_ fold change 2.8, adjusted P = 0.022; Supplementary Fig. 8; Supplementary Table 3). One gene among 248 tests, at a permulation P that would not survive correction across tests, is a lead for follow-up, not a finding.

### A perennial *Brachypodium* holds NRT2.3 at the level an annual reaches only under N deprivation

Sequence tests do not register differences in expression, so we compared root expression in annual and perennial *Brachypodium*, using public RNA-seq in which both were grown in the same experiments (Supplementary Table 21a). In one batch of greenhouse roots in fertilised soil (8 accessions of the annual *B. distachyon* and 6 of the perennial *B. sylvaticum*, three roots each; each species quantified on its own genome with transferred gene models), the species differed at 60% of 11,138 expressed genes (limma-voom with accession as a random block), but neither the orthologous uptake-state module nor the uptake and amino-acid-synthesis marker sets were shifted relative to all genes (camera P = 0.80, 0.97 and 0.23; Supplementary Table 21b). The NRT2.3 ortholog was the exception. Every perennial accession expressed it above every annual accession (accession medians 15-213 versus 0.4-6.1 TPM; log_2_ fold change 5.8, adjusted P = 3 × 10^−5^; exact Mann–Whitney P = 7 × 10^−4^; the 29th largest fold change of 11,138), and the AMT2;1 ortholog did the same at a smaller amplitude (17-32 versus 7.6-14 TPM; log_2_ 0.7, adjusted P = 6 × 10^−4^) (Fig. 7d, e; Supplementary Fig. 10a, b; Supplementary Table 21c). Both differences recurred in a second batch in which the perennial Ain-1 was compared with *B. distachyon* Bd21 and the annual *B. stacei* (NRT2.3 log_2_ 2.3 and 2.0, P = 0.03 and 0.04, with overlapping ranges; AMT2;1 log_2_ 1.5 and 1.0, P < 10^−8^).

The annual’s NRT2.3 is inducible. In *B. distachyon* Bd21-3 roots it rose from 1-2 TPM at 1.5 or 6 mM N to 21-32 TPM without N (log_2_ 3.5 for nitrate, P = 2 × 10^−5^; 2.7 for ammonium, P = 0.02; Supplementary Table 21d), in line with the strong down-regulation of NRT2 members at high nitrate reported by the producers of these data^57^, whereas *B. sylvaticum* roots grown with nitrate, ammonium or urea held 21-55 TPM (Fig. 7c). The *B. sylvaticum* NRT2.3 locus is intact on a large scaffold and only 69% identical to the NRT2.1 copies, which excludes cross-mapping, and *B. stacei*, whose gene models were transferred in the same way, shows the annual level. The NRT2.1 array and NAR2 could not be assessed because the *B. sylvaticum* assembly fragments them. With one perennial species, two comparisons spanning experiments and no N-starved *B. sylvaticum* sample, these data show that a perennial grass can run NRT2.3 at the level an annual congener reaches only under N deprivation, not that perenniality does so in general. Between species, this is the configuration that separates Barhoney from Fagerlin (Fig. 7a), on a gene whose coding sequence, copy number and proximal motifs carry no life-history signal.

### The bulk time course read through the atlas

We re-quantified the bulk time course (Barhoney, Fagerlin and the tetraploid Figgjo; 109 libraries retained of 114, which leaves the Barhoney 4 h full-N group with one library) on Kyuss_2.0. At 4 h the two data sets agreed on the direction of genotype differences (73% and 80% sign concordance for bulk-significant genes at full N and 25% less N; Spearman ρ = 0.32 and 0.65; Fig. 5e), but the bulk genes called as N-responsive at 4 h showed no correlation with the nuclear fold changes (ρ = −0.27 and 0.25).

Genotype differences at all 13 contrasts were enriched 1.7-3.9 fold for genes specific to the amino-acid-synthesis cell state (BH-adjusted Fisher P ≤ 0.03; Fig. 6a, b; Supplementary Table 15), whose module score was higher in Fagerlin in 11 of 13 groups (genotype term P = 1.3 × 10^−4^), whereas the PR/defence state scored higher in Barhoney (P = 0.012; Fig. 6c). Part of the whole-root genotype signal is therefore the signal of one cortical cell state, present from the starved baseline to 48 h. The uptake state, in contrast, was invisible in bulk: its module score showed no genotype, N or time effect (P = 0.60, 0.86 and 0.46), the NRT2 arrays were low in whole-root RNA (NRT2.3 pair ≤ 3 TPM; chromosome-4 and -6 arrays ≤ 40 TPM; Fig. 6d), and 73 of the 75 genes that respond to N in Fagerlin uptake cells showed no concordant change at any time (same sign 38-58%; 0-3 genes significant). A response confined to 17.5% of nuclei, one genotype and a low-abundance transporter family cannot be recovered from whole-root RNA.

### A public root atlas browser

The atlas is available at https://kovimallik.github.io/lolium-root-atlas/ (source at https://github.com/kovimallik/lolium-root-atlas, CC BY 4.0). The static page holds a 20,000-nucleus subsample of the embedding, the cell-state labels and the expression of all 35,145 genes detected in at least ten nuclei, stored as image files read back in the browser (59 MB in total). Users enter gene identifiers or a keyword and obtain expression on the UMAP, split by genotype, as violins per cell state or as a dot plot, each with a table of means and detection rates by genotype. A gene-list function annotates up to 200 identifiers with the cell state each gene marks, its specificity and its Barhoney-to-Fagerlin ratio, which is the question a breeder with candidates from association mapping asks of a root atlas. The subsample is for orientation, and statistics should use the full object (Data availability). To our knowledge this is the first root cell atlas for a forage grass, complementing genome-level resources such as ForageGrassBase for meadow fescue^58^; because the cell states are defined on genes conserved across Pooideae, orthologs from *Festuca*, *Dactylis*, *Phleum* and the cereals can be looked up once mapped to Kyuss_2.0 identifiers.

### Candidate markers for breeding

The variants that separate the genotypes fall into two classes of candidate marker (Supplementary Tables 5 and 12). DNA-level candidates are the six deletion-like genes, the 16 further genes without reads in the gene body, and the inactive LOC127308872 allele, each convertible to a PCR or KASP assay once the causal variant is known. Expression-level candidates are the 384 on/off genes and the paralog-specific NRT2 signature of the uptake cells (NRT2.3 and LOC127297370 2-fold higher in Barhoney under full N; LOC127308862 6-fold higher in Fagerlin), scorable by qPCR on root RNA with paralog-specific primers; because the NRT2 arrays as a whole are higher in Fagerlin in whole roots, family-level assays would not capture them.

## Discussion

The nitrogen response of the perennial ryegrass root has a cellular address. High-affinity nitrate uptake is concentrated in one cortical cell state, and the two genotypes run that state differently: the high-NUE genotype holds NRT2.3 at a higher baseline without induction, whereas the low-NUE genotype induces NRT2.3 and AMT2;1 and expands an induced sub-state. This explains much of what whole roots show. The uptake-cell response is confined to 17.5% of nuclei and almost entirely to one genotype, the thousands of genes by which the genotypes differ are enriched for the amino-acid-synthesis state and not for the uptake state, and the uptake state itself, with its low-abundance transporters, is invisible in whole-root RNA.

The cortical location (50% of uptake-state nuclei assigned to wheat cortex and 95% to maize cortex) differs from a wheat root-tip study that assigned N metabolism to the endodermis^24^ but agrees with the finding that outer root layers carry the transcriptional response to soil conditions^20^ and with the physiology of grass roots, in which epidermal and cortical cells respond electrically to nitrate uptake by the high-affinity system^59^ and the basal, mature part of the root, not the apex, carries nitrate acquisition^60^. The difference is most likely one of tissue, because we sampled whole roots, dominated by mature cortex, whereas root tips are enriched for meristem and young endodermis. Whole-root sampling also separated asparagine synthesis and amino-acid export into distinct states, an arrangement that places amino-acid exporters rather than a nitrate transporter in the stele, unlike the xylem-loading NRT2 of rice^38^.

We propose, as a hypothesis, that NUE differences in ryegrass arise from paralog deployment on a conserved toolkit. The uptake and assimilation families show no life-history change in copy number, rate or selection intensity beyond what random orthogroups show, yet the two genotypes use different NRT2 array members, one of them inactive in the high-NUE genotype, and the genes higher in the high-NUE genotype are modestly enriched for inducible-promoter elements. The same contrast between induced and uninduced transporters appears in a barley genotype pair, and between species the perennial *B. sylvaticum* holds NRT2.3 at the level that the annual *B. distachyon* reaches only under N deprivation (Fig. 7). The pattern resembles the “stress-prepared” transcriptomes of extremophyte Brassicaceae, whose roots constitutively express genes that sensitive relatives induce only under stress^25^: the high-NUE genotype behaves as nitrogen-prepared and the low-NUE genotype as nitrogen-responsive. It is also the pattern expected when divergence arises through cis-regulatory change in a redundant gene family^30–32^, the logic by which duplicated genes act as contingencies in crop engineering^33^. Known nitrate regulators (NLP2, PHR1-like GARPs, TGA bZIPs^17,18,39,40,61^) co-vary with the uptake module, which supports the identity of the uptake state, but their higher expression in the low-NUE genotype argues against a trans-acting explanation of the high-NUE baseline through these factors. Allelic imbalance at NRT2.3 in Fagerlin, but not in Barhoney, is consistent with a cis variant in the low-NUE genotype; an F1 between the two would test it directly.

To our knowledge, this is the first sequence-level comparison of NUE genes between annual and perennial grasses that judges candidate genes against random orthogroups and phylogenetic permulations. Across 44 trait-coded Pooideae genomes, copy number, relative rate and selection intensity on the uptake and assimilation families did not differ detectably with life history once phylogeny, ploidy, alignment size and the behaviour of random genes were taken into account. Significant shifts changed sign between contrasts, and 11 of 28 disappeared with one copy per species. The one robust test, relaxed constraint on the nitrate channel SLAH2 in wild perennials, coincides with higher endodermal SLAH2 expression in the high-NUE genotype, which we record as a lead. None of this implies that annuals and perennials handle nitrogen alike. The perennial N economy differs in where and when nitrogen moves: storage in stubble and roots and remobilisation for regrowth after each cut and over winter, instead of a single transfer to grain (Supplementary Fig. 9). Such differences can arise through regulation of conserved genes, through programmes such as senescence and storage-tissue identity that lie outside the N toolkit, and through different changes in each of the 10-17 origins of annuality, none of which per-gene tests of coding constraint, rate or copy number can detect. The *Brachypodium* comparison is a first test of the regulatory route, in one genus and without an N-starved perennial sample; repeating it across independent origins of perenniality and across a defoliation and regrowth cycle would test it directly.

The design has limits. Each NUE class is represented by one clonal genotype, so genotype and NUE class cannot be separated. The two plants also differed in growth state (the low-NUE genotype had more dividing and ribosome-rich nuclei, the high-NUE genotype more secondary-wall and phenylpropanoid expression; Supplementary Tables 9 and 11), which contributes to the genome-wide differences and could contribute to baseline uptake-gene expression, though not to the inactive paralog or the within-genotype N response. The treatment was a single 4 h resupply contrast after one week of starvation, so “constitutive” and “inducible” describe this contrast only, and nuclear RNA under-represents stable cytoplasmic transcripts. Association of the candidate variants with NUE across the field panel, in situ hybridisation in additional genotypes (candidate probe regions are listed in Supplementary Table 10) and loss of function of the NRT2.3 pair, for which genome editing is being developed in forage grasses^62^, would address these limits.

For breeding, the relevant variation is visible in roots under sufficient N, as paralog-specific NRT2 expression in the uptake cells and as presence/absence and on/off variants that standard SNP genotyping does not capture. Perennial ryegrass is bred as synthetic populations and increasingly by genomic selection^11^, and the uptake-state markers, once validated in the field panel, offer a concrete route to pastures that need less nitrogen. Because the uptake state is built from genes conserved across Pooideae, the same markers can be sought in other forage grasses.

## Methods

### Plant material and nitrogen treatment

Barhoney (high field NUE) and Fagerlin (low field NUE), both diploid, were selected from a field trial of 40 cultivars grown at Ås and Hamar, Norway, under three N regimes with three replicates, in which NUE was calculated as harvested yield per unit of N applied. Clonal tillers of one plant per genotype were acclimated in aerated modified Hoagland solution^63^ at full N (14.0 mM nitrate), starved of N for 7 days, then transferred at the same time of day to solution with full N (control; 14.0 mM nitrate) or 25% less N (10.5 mM nitrate); the bulk time course used the same system with a third level, 15% less N (11.9 mM nitrate). Whole root systems of three plants per group were harvested 4 h later, rinsed and snap-frozen in liquid nitrogen.

### Nuclei isolation, libraries and sequencing

Nuclei were isolated following the Novogene plant single-nucleus protocol and loaded on 10x Chromium GEM-X Single Cell 3′ v4 targeting 10,000 nuclei per library^64^; two libraries (L07, L10) were processed as a pilot and ten in a second batch. Libraries were sequenced PE150 on Illumina NovaSeq X to 91-121 Gb (305-405 million read pairs) per library.

### Primary processing

Reads were processed with Cell Ranger 8.0.1^64^ (include-introns) against the NCBI RefSeq Kyuss_2.0 assembly and annotation (GCF_019359855.2, RS_2025_02; 56,405 genes)^34,35^. Mapping rates were 33-65%; re-alignment of 200,000 reads from each of four libraries with minimap2^65^ showed that ∼45% of reads were rRNA (multi-mapping to rDNA scaffolds) and <5% were unmapped. Cell calling was Cell Ranger’s (EmptyDrops-based^66^); nuclei retained by the provider’s Seurat/DoubletFinder^67^ filters (88,323 nuclei, 28 provider clusters) formed the input set for our processing, and the provider clusters were used for ambient estimation, label transfer and the whole-root comparisons noted below.

### Ambient correction, doublets and integration

Ambient RNA was removed with SoupX 1.6.2 (autoEstCont per library; ρ = 0.05-0.15)^68^ and doublets with scDblFinder 1.23.4 on corrected counts (9-17% removed)^69^; nuclei with <300 genes or >10% rRNA counts were excluded, leaving 76,684. In Scanpy 1.11^70^ counts were log-normalised, 3,000 highly variable genes selected batch-aware across libraries (Seurat flavour, batch = library), 50 principal components computed and integrated with Harmony over library^71^, followed by a k-nearest-neighbour graph (k = 20), UMAP^72^ and Leiden clustering (resolution 1.0; 35 clusters)^73^.

### Cell-type annotation and label transfer

Clusters were annotated from marker enrichment (4,458 marker × cell-type records covering 2,938 genes, derived from published atlases^21,22,74,75^ and mapped to Kyuss_2.0), RefSeq gene descriptions, and label transfer run on the 28 provider clusters: scANVI^76^ and scanpy ingest trained on the wheat root atlas GSE270342^23^ (authors’ labels; 14,414 Lolium-space genes built by summing the one to three wheat homoeologs orthologous to each Lolium gene, OrthoFinder^44^), plus Spearman correlation of cluster pseudobulks to wheat cell-type signatures; a label was accepted when two of these three methods agreed (16 of 28 clusters), and scANVI trained on the maize atlas GSE183171^21^ (8,830 one-to-one orthologs) was used as an independent coarse-level check. The 35 Leiden clusters of the cleaned object inherited labels by majority overlap with the provider clusters and were then reviewed against markers (Supplementary Table 2). The whole-root pseudobulk comparison, replicate correlations, marker enrichment, label transfer, leave-one-library-out and pilot-batch sensitivity analyses and the GO enrichment were computed on the provider object (88,323 nuclei); all other results use the cleaned object (76,684 nuclei).

### Differential expression and composition

Counts were summed per library within each cluster or cell-type group (pseudobulk; ≥20 nuclei per library and ≥10 of the 12 pseudobulks present; six small clusters were skipped)^77^ and analysed with DESeq2 1.46^78^ using ∼pilot + group (group = genotype × treatment) with contrasts for the N response within genotype and the genotype difference within treatment, and ∼pilot + genotype × treatment for the interaction; padj < 0.05 (Benjamini–Hochberg^79^). Robustness was assessed by leave-one-library-out and batch-covariate sensitivity analyses. Composition was tested propeller-style^80^ (limma^81^ on logit proportions, ∼genotype + treatment + pilot; Supplementary Table 11). Gene Ontology and KEGG enrichment used clusterProfiler^82^ with a Lolium annotation built by orthology (Supplementary Table 9).

### Sub-clustering and regulator inference

Uptake nuclei were re-clustered on the Harmony embedding (Leiden 0.6). Transcription factors (1,485 by RefSeq family keywords, of which 333 were detected in ≥2% of uptake nuclei and tested) were correlated (Spearman) with an uptake-module score (23 panel genes) and an induced-gene score on 200-nucleus within-library metacells, with bootstrap confidence intervals and depth-partial and within-genotype controls; no motif databases exist for *Lolium*, so inference is expression-based.

### Paralogs, promoters and presence/absence

NRT2 paralog identity was computed by blastn^83^ of RefSeq transcripts. Promoter motifs (1 kb and 500 bp upstream, strand-aware, Kyuss_2.0) were tested by Fisher’s exact test against expression and GC-matched backgrounds, Cochran–Mantel–Haenszel stratification and 1,000 matched permutations. On/off genes were defined on whole-root pseudobulk (≥2 CPM in all six libraries of one genotype and ≤0.05 CPM in all six of the other) and classified from unique-read (MAPQ 255) coverage of gene bodies and ±2 kb flanks (samtools^84^). Allele-specific expression used bcftools^84^ mpileup/call (MAPQ ≥ 20) over expressed gene bodies per genotype (six libraries pooled; QUAL ≥ 30, DP ≥ 30, pooled allele fraction 0.2-0.8), per-library allelic depths at those sites, and per-cell-type allelic depths from BAMs split by cluster barcodes. Gene-level imbalance was estimated with MBASED 1.40^41^ in one-sample unphased mode on counts pooled over the six libraries of a genotype, with the genome-wide pooled reference-allele fraction of that genotype (0.507 Barhoney, 0.473 Fagerlin) as the per-site no-ASE expectation, a beta-binomial dispersion estimated by the method of moments from per-library allele fractions at sites with ≥30 reads in every library (ρ = 0.006 and 0.021), 10^6^ simulations per gene for the focal genes and 10^4^ genome-wide, and the authors’ call rule (BH-adjusted P_ASE_ < 0.05 and major-allele frequency ≥ 0.7). Library-level reproducibility, which MBASED does not test, was checked with a leave-one-library-out orientation (each site oriented by the pooled counts of the other five libraries; the held-out library’s fraction has expectation 0.5 under the null; sites ≥150 bp apart) and is reported beside the MBASED estimate (Supplementary Table 13); applied to counts simulated under the null at the observed depths, this check produced no false call. For the 24 focal genes, sites whose 31-mer carrying the alternative allele occurs elsewhere in the reference were removed as paralog-explainable (blastn-short, 100% identity). Because heterozygous calls in multi-copy families can also arise from copies absent from the reference, results for the NRT2 arrays are reported descriptively. RNA velocity used spliced/unspliced molecule counts derived from the Cell Ranger region tags of each read (unspliced = molecule with any intronic read) and scVelo 0.3 (stochastic and dynamical models on the Harmony neighbour graph; latent time per nucleus summarised by cell state, genotype and treatment)^85,86^; because only 10.8% of molecules were unspliced, velocity is reported descriptively.

### Web viewer

From the integrated object, 20,000 nuclei were sampled at random (seed 0); UMAP coordinates (16-bit), cell-state and library indices (8-bit) and, for every gene detected in ≥ 10 nuclei, log1p counts per 10,000 scaled to the gene’s 99th percentile among expressing nuclei (8-bit) were written as PNG images (128 genes × 20,000 nuclei per file) and a JSON gene index with RefSeq descriptions. The page (plain HTML and JavaScript, no dependencies) reads the images through a canvas and draws UMAP, genotype-split, violin and dot-plot views and per-state summaries client-side; it is served by GitHub Pages from the repository.

### Bulk RNA-seq

114 libraries (three cultivars, three N levels, five time points; 19.9-37.2 million read pairs each, median 26.5 million) were trimmed with fastp^87^ and quantified with Salmon 1.10 (gcBias, selective alignment, genome decoys)^88^ against the Kyuss_2.0 RS_2025_02 transcriptome (and, for comparison with earlier results, Kyuss v1^34^); five libraries with low mapping rates were excluded, leaving 109. Gene counts were obtained with tximport^89^ and analysed with DESeq2 (design ∼group; genes with ≥5 counts in ≥3 libraries) using the 4 h Barhoney/Fagerlin libraries (control versus 25% less N; Barhoney control n = 1). For the time course, the 70 Barhoney and Fagerlin libraries (genes with ≥10 counts in ≥3 libraries; 35,617 genes) were modelled as ∼group (group = genotype × N level × time) with contrasts for time versus 0 h at full N, for 15% and 25% less N versus full N at each time, and for Barhoney versus Fagerlin at each N level and time; groups containing one library are reported but flagged. Cell-state specificity of each gene was computed from the snRNA-seq label-level pseudobulks (tau index^90^ over the 15 cell states; specific if tau ≥ 0.6 and ≥5 CPM), the 50 most specific genes per state were scored in each bulk library as the mean z-score of log2(TPM + 1), module scores were tested with linear models (score ∼ genotype + N level + time; sequential analysis-of-variance P values), and the enrichment of state-specific genes among each DEG set relative to all expressed genes was tested with one-sided Fisher’s exact tests (BH-adjusted).

### Public root N-response data

Root RNA-seq runs of barley PRJNA896249^42^ (W26, W20; CK, LN at 3 and 18 d; LN and RN at 21 d; 36 runs), Brachypodium PRJNA1348082^57^ (nitrate and ammonium series) and wheat PRJEB54622, PRJNA1082516 and PRJDB38906 (high versus low N; nitrate induction) were downloaded from ENA and quantified with salmon (--validateMappings --gcBias) against the Ensembl Plants cDNA sets (MorexV3, Brachypodium v3.0, IWGSC), transcripts summed per gene. Orthologs of the 50-gene uptake-state module (39-44 with orthologs per species), of the amino-acid-synthesis module and of the core N-gene panel were assigned by DIAMOND^91^ best hit (very-sensitive, e < 10^−10^, query coverage ≥ 60%) from the longest Kyuss_2.0 protein per gene, with reciprocal best hits flagged and family members retained where the reverse best hit fell in the same family. Per genotype, module and gene responses were summarised as log_2_ fold changes of mean TPM (+1) with Welch tests on per-library values and the genome-wide fold-change distribution as reference; the Lolium bulk series was treated identically (Supplementary Table 20).

### Annual and perennial *Brachypodium* roots

Root RNA-seq runs of the Joint Genome Institute *Brachypodium* series (Supplementary Table 21a) were downloaded from NCBI SRA (first 10 million spots per run). Batch R18 (“Biomass Allocation Gene Expression Profiling”) contains 8 accessions of the annual *B. distachyon* and 6 of the perennial *B. sylvaticum*, three roots each, grown together in a greenhouse (25/20 °C, well-fertilised and well-watered soil); batch R19 (“Roots and Leaves Gene Expression Profiling”) contains *B. distachyon* Bd21, the annual *B. stacei* ABR114 and *B. sylvaticum* Ain-1 under the same description; *B. sylvaticum* Ain-1 roots grown in a growth chamber with nitrate, ammonium or urea as nitrogen source are from Gene Atlas plate 15 (PRJNA411649)^92^, whose records do not state concentrations. A *B. distachyon* Bd21-3 nitrogen-source series of the same programme was shoot tissue by its photosynthesis transcripts (9-20% of TPM in RBCS and LHCB genes, against ≤ 1.4% in every root sample) and was not used. Each species was quantified with salmon^88^ (--validateMappings --gcBias) on its own transcriptome: *B. distachyon* RefSeq (GCF_000005505.3; protein-coding genes, organelle genes removed) and, for *B. sylvaticum* (GCA_040967415.1) and *B. stacei* (GCA_036872525.1), which have no public annotation, the same models transferred with Liftoff 1.6.3 (-polish)^93^, so that a gene identifier names the same locus in the three species; gene-level counts were imported with tximport (lengthScaledTPM)^89^. Runs with < 50% of reads mapped (three) were excluded, technical re-runs of one sample were summed, and genes were restricted to those transferred with ≥ 90% coverage and a valid open reading frame. In R18, genes with ≥ 1 CPM in ≥ 3 libraries (11,138) were tested with limma-voom^81,94^ (TMM normalisation; life history as the factor; accession as a random block via duplicateCorrelation, so that accessions rather than plants are the replicates); in R19 (25 libraries, 9,194 genes) the perennial was contrasted with each annual. Lolium genes were mapped to *B. distachyon* RefSeq genes by DIAMOND as above (genome-wide reciprocal best hits; best hits for the modules), and N-family members whose *B. distachyon* product contradicted the family were removed; gene sets were tested with camera^95^ (inter-gene correlation 0.01) on accession means, and accession medians of single genes with an exact Mann–Whitney test. The *B. distachyon* NRT2.3 ortholog in the Bd21-3 nitrate and ammonium series (PRJNA1348082^57^; 9-day-old roots) was read from the quantification of the preceding paragraph (BRADI_2g47640, the same locus as LOC100835409), with Welch tests of 0 versus 6 mM on log_2_(TPM + 1).

### Comparative genomics

Genomes (53 assemblies, 48 Pooideae core; Supplementary Table 6) were obtained from NCBI, and life history, cultivation status, ploidy level and assembly level were curated from the botanical literature, chromosome counts and assembly reports (the haplotype-resolved *Hordeum bulbosum* assembly was reduced to one haplotype; the 44 core genomes with high-confidence life-history coding enter the contrasts). Members of 25 N-use and associated families were mined with miniprot^43^ from seed proteins of *Arabidopsis*, rice, *Brachypodium* and barley (the nitrate-reductase panel included the molybdenum-cofactor genes CNX2 and CNX3, which form their own orthogroups), overlapping models were collapsed to loci, and only intact loci (no frameshift or internal stop, ≤5% ambiguous residues, length 0.5-1.6× the seed median, ≥60% query coverage, ≥35% identity) were counted (24 families in the copy-number analysis). Copy number was modelled by phylogenetic generalised least squares (caper^96^; Pagel’s λ by maximum likelihood^51^) on the species tree made ultrametric with ape::chronos^97^, as log(1 + copies) ∼ life history + ploidy level + assembly level over the 42 genomes with high-confidence life history and curated ploidy, repeated on diploids only (the chromosome-level-only subset did not yield a converged fit), and by sign tests of the annual–perennial difference in mean copy number within the five genera containing both life histories. Orthogroups were inferred with OrthoFinder^44^ guided by the species tree, which was built with IQ-TREE 2 from a supermatrix of single-copy orthologues (87,033 amino-acid sites; ModelFinder; 1,000 ultrafast bootstraps)^45–47^. Hierarchical orthogroups with ≥12 core species (200 of 214) were filtered for sequence length (0.7-1.15× the orthogroup median), aligned at the protein level with MAFFT^98^ (L-INS-i for ≤150 sequences, otherwise --auto) and back-translated to codons, trimmed to codon columns with ≥50% occupancy (sequences with <50% non-gap positions and alignments shorter than 100 codons dropped; 198 orthogroups with gene trees), and gene trees were inferred with IQ-TREE 2; tips on branches longer than 0.5 substitutions per site were removed (two). For each contrast, tips were labelled Test or Reference from their species and internal branches by parsimony from their descendants, nodes with mixed descendants being left unclassified; contrasts with fewer than three species on either side or fewer than six tips were not tested, and alignments with more than 120 sequences were subsampled to two copies per species. Selection intensity was tested with RELAX^53^ (synonymous rate variation, ten starting points) and trait-associated episodic selection with BUSTED-PH^54,55^, in HyPhy 2.5.96^99^, with Benjamini–Hochberg correction^79^ within each contrast; k estimates below 0.1 or above 10, or fits with an ω class above 100 carrying less than 1% weight, were treated as non-interpretable, and a BUSTED-PH call required foreground P ≤ 0.05, background P > 0.05 and a significant difference between the two distributions. The empirical background was 100 random single-copy orthogroups, drawn from 1,971 single-copy markers across the core genomes, run through the identical labelling and tests and compared by Fisher’s exact tests on hit rates, by logistic regression of a shift at P < 0.05 on gene set and log number of sequences per test, per contrast and overall (because the multi-copy N-use orthogroups yield larger alignments than single-copy controls; Supplementary Table 7e), and by Mann–Whitney tests on k; the long-branch filter removed the marker models of the large genomes that lay on saturated branches (15-63% of the markers in the 18 genomes ≥4 Gb, ≤1% in smaller genomes; Supplementary Note 1), so the control alignments under-represent the large genomes, which the logistic adjustment on alignment size partly absorbs. Relative evolutionary rates were computed with RERconverge 0.3.0^52^: for each of the 198 orthogroups the sequence with the fewest gaps per species was taken from the MAFFT protein alignment, branch lengths were re-estimated on the species-tree topology pruned to the species present (phangorn^100^, LG model), relative rates were obtained with getAllResiduals (square-root transform, weighted, species of the contrast only) and correlated with the binary trait (Test species as foreground, clade = all, matching the parsimony labelling above) with correlateWithBinaryPhenotype, BH-corrected within the core set per contrast, and tested against 500 binary permulations (getPermsBinary, complete-case mode) on the pruned species tree^101^; hit rates were compared with the 148 non-core orthogroups by Fisher’s exact test. A project-specific root-to-tip measure on the multi-copy gene trees is reported in Supplementary Note 1 as a sensitivity analysis only; the single-copy marker set was not used as the rate baseline for the reason given above. Promoter elements were scored in the 1-kb region upstream of the start codon of every kept model (samtools faidx, strand-aware; models with < 900 bp or > 10% N excluded) with the IUPAC consensus set of the snRNA-seq promoter analysis on both strands, as presence per model, averaged per species and orthogroup, and tested per contrast by the same Brownian-motion permulation scheme (1,000 permulations) against the 150 non-core orthogroups, with GC content tested alongside. Nominally significant RELAX tests were refitted twice more (ten starting points each) and the best-likelihood fit is reported with the replicate agreement; the same tests were run through aBSREL^102^ and Contrast-FEL^103^, repeated with one copy per species (longest sequence; internal branches re-labelled), and tested against species-tree Brownian-motion permulations^101^ (null foregrounds of the same species count; two-sided P on |ln k|; 50 permulations per test, of which 42-49 gave interpretable null fits, and 89 for the retained SLAH2 test). The design and the full results are given in Supplementary Note 1.

### Statistics and reproducibility

All analysis code (R and Python, numbered scripts) and environment specifications are available from the corresponding author. Sample sizes, tests and corrections are stated with each result.

## Data availability

Bulk RNA-seq data, raw snRNA-seq reads, Cell Ranger matrices and the processed atlas object (AnnData with raw counts, normalised expression, cell-state annotations and embeddings) will be deposited in an open-access public repository. An interactive web viewer of the atlas (all detected genes, cell states, genotypes) is at https://kovimallik.github.io/lolium-root-atlas/. Genome assemblies are listed in Supplementary Table 6. The wheat and maize reference atlases are GSE270342^23^ and GSE183171^21^.

## Code availability

Analysis scripts are available from the corresponding author. The web viewer and its build script are at https://github.com/kovimallik/lolium-root-atlas.

## Acknowledgements

The *Brachypodium* root RNA-seq data analysed here were produced by the US Department of Energy Joint Genome Institute (JGI) Community Science Program and the JGI Plant Gene Atlas; we thank the JGI and the principal investigators of these projects. This work is part of DLT-Farming, NitrogenEdit, Soil2Milk projects funded by FFL/JA, Landbruksdirektorate, and Norges Forskningsråd and NMBU for PhD grant to Precious, Novogene for library preparation and sequencing; NMBU for computing resources.

## Supplementary information

Supplementary Figures 1-10, legends of Supplementary Tables 1-21 and Supplementary Note 1 are provided in the Supplementary Information file; Supplementary Tables 1-21 are provided as an Excel workbook, one table per sheet.

